# Multi-omic regulatory networks capture downstream effects of kinase inhibition in *Mycobacterium tuberculosis*

**DOI:** 10.1101/584177

**Authors:** Albert T. Young, Xavier Carette, Michaela Helmel, Hanno Steen, Robert N Husson, John Quackenbush, John Platig

**Affiliations:** School of Medicine, University of California, San Francisco, California, USA; Division of Infectious Diseases, Boston Children’s Hospital, Boston, Massachusetts, USA; Harvard Medical School, Boston, Massachusetts, USA; Department of Pathology, Boston Children’s Hospital, Boston, Massachusetts, USA; Department of Biostatistics, Harvard T.H. Chan School of Public Health, Boston, Massachusetts, USA; Department of Data Sciences, Dana-Farber Cancer Institute, Boston, Massachusetts, USA; Channing Division of Network Medicine, Brigham and Women’s Hospital, Boston, Massachusetts, USA

## Abstract

The ability of *Mycobacterium tuberculosis (Mtb)* to adapt to diverse stresses in its host environment is crucial for pathogenesis. Two essential *Mtb* serine/threonine protein kinases, PknA and PknB, regulate cell growth in response to environmental stimuli, but little is known about their downstream effects. By combining RNA-Seq data, following treatment with either a PknA/PknB inhibitor or an inactive control, with publicly available ChIP-Seq and protein-protein interaction data, we show that the *Mtb* transcription factor (TF) regulatory network propagates the effects of kinase inhibition and leads to widespread changes in regulatory programs involved in cell wall integrity, stress response, and energy production, among others. We also observe that changes in TF regulatory activity correlate with kinase-specific phosphorylation of those TFs. In addition to characterizing the downstream regulatory effects of PknA/PknB inhibition, this demonstrates the need for regulatory network approaches that can incorporate signal-driven transcription factor modifications.

## Introduction

*Mycobacterium tuberculosis* (Mtb) remains one of the world’s deadliest pathogens, with 10.4 million people falling ill with tuberculosis disease (TB) and over 1.6 million TB deaths in 2016 (*1*). The emergence of multidrug-resistant and extensively drug-resistant *Mtb* strains threatens to undermine global TB control efforts.

Much of *Mtb*’s success as a pathogen can be attributed to its ability to adapt to diverse environmental stresses encountered during the course of chronic infection (*2*). Protein kinases anchored on the mycobacterial cytoplasmic membrane are critical for responding to environmental stimuli and transducing signals to various cellular processes (*3*). The essential *Mtb* serine/threonine protein kinases (STPKs) PknA and PknB are excellent targets to characterize in the context of future drug development, as they regulate several processes required for cell growth and division, including the biosynthesis of essential components of the cell envelope (peptidoglycan, mycolic acids, and other cell wall lipids and carbohydrates) (*4*). For example, cells in which PknA or PknB gene expression was inhibited displayed an abnormal shape, indicating the two kinases are key regulators of cell division and cell shape in *Mtb* (*5*). However, our understanding of the downstream transcriptional pathways by which PknA and PknB regulate these and other cellular processes is limited, and the basis of their essentiality is unknown.

To measure the downstream effects of PknA/PknB signaling, we used a potent small molecule inhibitor of both PknA and PknB along with an inactive negative control, then collected multiple omics data types as detailed in (*6*). To follow the propagation of this signaling perturbation, we sought to identify differences in transcription factor (TF) regulation in the inhibitor- and control-treated gene expression programs by reconstructing TF regulatory networks in the active (i.e. inhibitor) and control conditions. While condition-specific binding information for all TFs was not available, we combined recently generated ChIP-seq data for 143 TFs (as defined in (*7*)) in *Mtb* and protein-protein interaction data (*8*) with RNA-Seq data from inhibitor or control treated samples using the TF regulatory network reconstruction algorithm PANDA (*9*) (see Figure 1).

**Figure 1:**
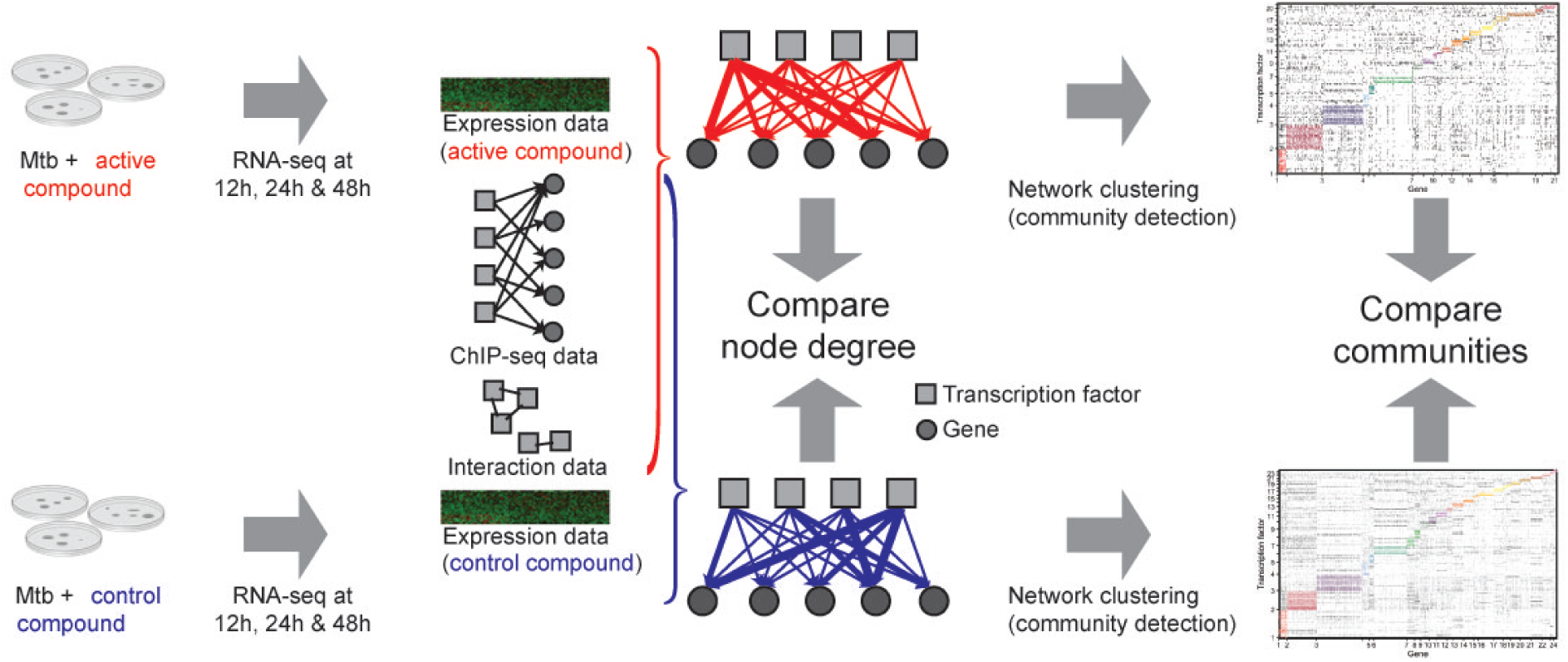
Summary of the experimental design and network comparisons. Using a message passing framework (*9*) (see Methods) we integrated ChIP-seq and protein-protein interaction (PPI) data with condition-specific gene expression data to build kinase-inhibitor and control gene regulatory networks. Network edges connect TFs to genes and are weighted to reflect the confidence (in z-score units) of a regulatory relationship based on the concordance between omics data types. The TF node degree is defined as the sum of all weights for edges emanating from the TF, and gene degree is the sum of all weights for edges terminating at the gene. The inhibitor and control networks were clustered into groups containing both TFs and genes using bipartite community detection (*24, 27*) as detailed in the Methods.

By comparing network topologies of these inhibitor and control networks, we first identify TFs that change their regulation in response to PknA/PknB inhibition as measured by change in network outdegree (sum of outbound links). We also show that change in network outdegree is predictive of change in phosphorylation status of the TF after PknA/PknB inhibition, suggesting that our network approach is modeling the downstream effects of the kinase inhibition. Second, we show that genes that are differentially regulated as measured by their change in indegree (sum of inbound links) are enriched for multiple functions, including mycobactin synthesis, with additional validation that mycobactin levels are indeed changed upon PknA/PknB inhibition (*6*). Third, we demonstrate that network “communities” (modules) in the inhibitor-treated network show condition-specific functional enrichment, which are validated by follow-up experiments.

## Main Text

### Change in TF outdegree is predictive of change in TF phosphorylation

To test whether the signaling perturbation of PknA/PknB inhibition was detected in our networks, we used phosphoproteomic data collected from samples treated with the PknA/PknB inhibitor or control and calculated log2 fold change values to determine phosphorylation status of the TFs in our networks. Peptides for 14 TFs were detected, 11 of which were differentially phosphorylated at an adjusted P-value < 0.05. We then compared the change in outdegree (defined as the sum of all out-going edge weights) between the inhibitor and control networks for each TF to the change in phosphorylation (see Figure 2) and find that these are significantly correlated (Spearman’s rho of 0.618, *P* = 0.0213). MtrA is one such TF. It has the 13th (out of 143) largest change in outdegree between the inhibitor and control networks. MtrA is also ranked third in differential phosphorylation among the 14 TFs measured (see Fig. 2). We also recently demonstrated that phosphorylation of MtrA inhibits DNA binding to the FbpB promoter (*6*).

**Figure 2:**
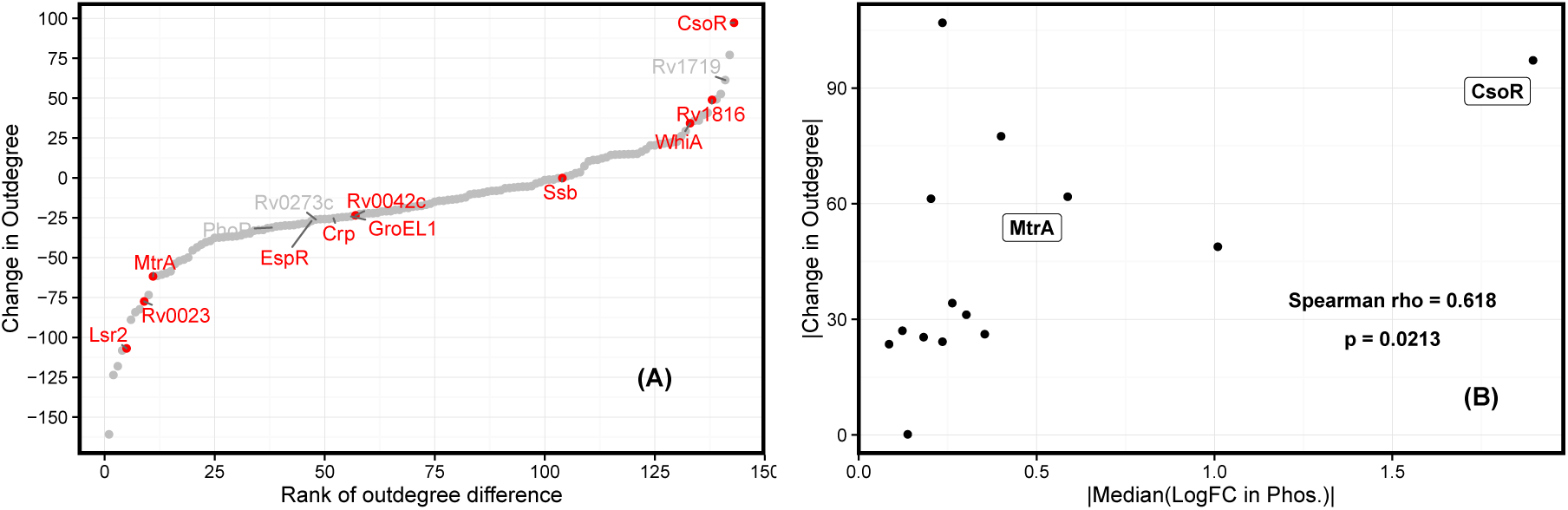
Change in transcription factor phosphorylation is correlated with change in outdegree. (A) The outdegree difference (*y*-axis) and rank of outdegree difference (*x*-axis) are shown for the 143 transcription factors included in the regulatory network model. Of those, 14 TFs had detected phosphopeptides and 11 were differentially phosphorylated (adj. P < 0.05, shown in red). (B) The magnitude of change in TF outdegree between active and control kinase inhibitor networks correlates with change in phosphorylation (Spearman correlation, *ρ*_*s*_ = 0.618 and *P* = 0.0213). The median log2 fold change was used when multiple phosphopeptides were detected for the same protein.

**Figure 3:**
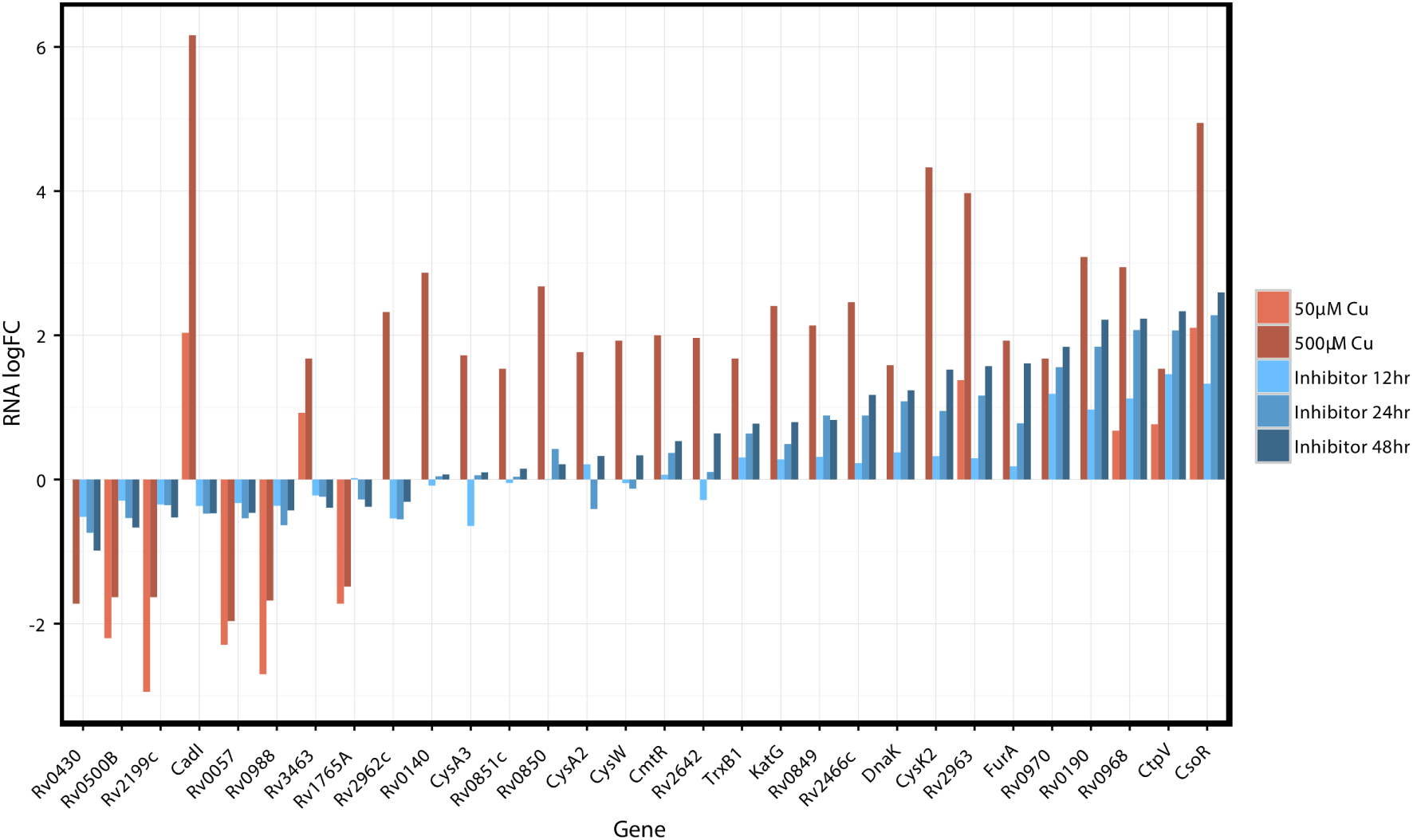
The transcriptional response of the CsoR regulon to PknA/PknB inhibition mirrors that of copper exposure. Log2 fold changes in RNA for genes in the CsoR regulon after treatment with the PknA/PknB inhibitor are shown in blue, changes in RNA after treatment with copper are shown in red (copper data from (*28*)). The copper sensitive operon repressor (CsoR) showed the greatest change in phosophorylation and increase in outdegree upon PknA/PknB inhibition (see Fig. 2).

This discovery suggests that changes in transcriptional regulation in response to PknA/PknB inhibition can be partly attributed to differential phosphorylation of TFs. In particular, CsoR, the most positively differentially targeting TF, is significantly differentially phosphorylated with a log2 fold change of −1.89. That the majority (8/11) of differentially phosphorylated TFs in the network have decreased outdegree in the inhibitor versus control network likely reflects the disruption of gene regulatory programs in response to PknA/PknB inhibition (i.e. genes in TF regulons have less correlated expression).

### Comparing networks uncovers PknA/PknB-specific targeting patterns of TFs

The PknA/PknB-specific TFs (Table 1 and S1 Table) are involved in processes we expect to be differentially regulated in the context of PknA/PknB inhibition. To assess this in the network context, we define “differential targeting” to mean the change in a TF’s outdegree between active control and kinase inhibitor networks. Rv0081, the most differentially targeting TF, is a regulatory hub in the context of hypoxia, a condition which induces a stress response with similarities to PknA/PknB inhibition (*10*). Other top differentially targeting TFs include the response regulator TrcR, which activates its own coding gene expression and represses Rv1057, a *β*-propeller protein gene whose expression is also mediated by SigE (*11*). Lsr2 is a global transcriptional regulator that may be responsible for many cell wall functions and is required for adaptation to changing oxygen levels (*12, 13*). CsoR, the TF with the greatest increase in targeting in kinase inhibitor treated cells, is a copper sensing transcriptional regulator that may promote *Mtb* survival by mediating a response to copper toxicity (*14*). KstR, a transcriptional repressor, controls a number of genes involved in cholesterol and fatty acid catabolism (*15*). Thus, each of the top 10 PknA/PknB-specific TFs with known function is involved in either signal transduction, cell wall function, or lipid metabolism–processes which PknA and PknB regulate. Given this, we propose Rv0678, Rv0324, Rv0465c, Rv1985c, and Rv0023, which have unknown functions, to be candidate downstream regulators most affected by PknA/PknB inhibition as determined by change in outdegree.

**Table 1:** Top ten transcription factors ranked by absolute change in outdegree difference between control and inhibitor networks. P-values are calculated from permutations as outlined in the Methods.

| RV | Degree Diff. | P-value | Gene | Product |
| --- | --- | --- | --- | --- |
| Rv0081 | -160.8 | 0.001 |  | transcriptional regulator, ArsR family |
| Rv1033c | -123.6 | 0.001 | TrcR | DNA-binding response regulator TrcR |
| Rv0678 | -118.1 | 0.001 |  | Conserved protein |
| Rv0324 | -108.2 | 0.001 |  | Transcriptional regulator, ArsR family |
| Rv3597c | -106.9 | 0.001 | Lsr2 | Histone protein Lsr2 |
| Rv0967 | 97.2 | 0.021 | CsoR | Copper-sensitive operon repressor CsoR |
| Rv0465c | -89.0 | 0.005 |  | Transcriptional regulator, XRE family |
| Rv1985c | -84.2 | 0.001 |  | HTH-type transcriptional regulator |
| Rv3574 | -82.4 | 0.005 | KstR | Transcriptional regulator kstR (Rv3574), TetR family |
| Rv0023 | -77.5 | 0.002 |  | Transcriptional regulatory protein |

Among the genes that are “differentially targeted”, which we define as the difference in gene indegree between control and inhibitor networks (Table 2 and S2 Table), we discovered many genes with interesting functions that may play important roles in responding to PknA/PknB inhibition. VapB30 and VapB40 are antitoxins in the VapBC (Virulence associated protein) family of toxin-antitoxin systems that regulate translation in response to diverse environmental stresses (*16*), and both VapB30 and VapB40 have increased gene expression and increased indegree in kinase inhibitor treated cells. CysD is involved in sulfur metabolism, which may be important in the virulence, antibiotic resistance, and antioxidant defense mechanisms of *Mtb* (*17*). ArgC, N-acetyl-gamma-glutamyl-phosphate reductase, is involved in arginine biosynthesis, an essential metabolic function for cellular growth and a pathway that is required for virulence (*18, 19*).

**Table 2:** Top ten genes ranked by absolute indegree difference between inhibitor and control networks. P-values are calculated from permutations (see Methods).

| RV | Degree Diff. | P-value | Gene | Product |
| --- | --- | --- | --- | --- |
| Rv2913c | 64.6 | 0.001 |  | N-acyl-D-glutamate amidohydrolase |
| Rv0623 | 63.2 | 0.001 | VapB30 | Possible antitoxin VapB30 |
| Rv3182 | 47.5 | 0.001 |  | Conserved hypothetical protein |
| Rv1285 | 46.8 | 0.001 | CysD | Sulfate adenylyltransferase subunit 2 (EC 2.7.7.4) |
| Rv1652 | 41.2 | 0.001 | ArgC | N-acetyl-gamma-glutamyl-phosphate reductase (EC 1.2.1.38) |
| Rv2282c | -37.9 | 0.001 | CysB | Cys regulon transcriptional activator CysB |
| Rv3453 | 37.4 | 0.001 |  | PROBABLE CONSERVED TRANSMEMBRANE PROTEIN |
| Rv1219c | 37.3 | 0.001 |  | Transcriptional regulator, TetR family |
| Rv2595 | 36.2 | 0.001 | VapB40 | Possible antitoxin VapB40 |
| Rv1693 | -35.1 | 0.001 |  | Conserved hypothetical protein |

Using gene set enrichment analysis (GSEA) for difference in indegree, we identified PknA/PknB-specific functional categories (S3 Table). “Mycobactin biosynthesis,” “phosphopantetheine binding,” and “ESX-1 LOCUS” show increased targeting in the inhibitor condition, and “Oxidative phosphorylation,” “quinone binding,” and “NADH dehydrogenase” show decreased targeting. Mycobactins are essential for iron acquisition within the host environment (*20*), and in related work (Figure 4 of (*6*)), we found that mycobactin levels were increased 48 hours after inhibition of PknA/PknB. “Phosphopantetheine binding” consists of 15 genes, including polyketide synthases crucial for fatty acid synthesis (*21*) and enzymes involved in mycobactin synthesis (*22*). The ESX-1 system is a specialized secretion system required for virulence (*23*). Oxidative phosphorylation, quinone binding, and NADH dehydrogenase genes are less targeted after PknA/PknB inhibition, supporting their role in mediating a compensatory biological response in the form of lowered energy expenditure and growth arrest.

**Figure 4:**
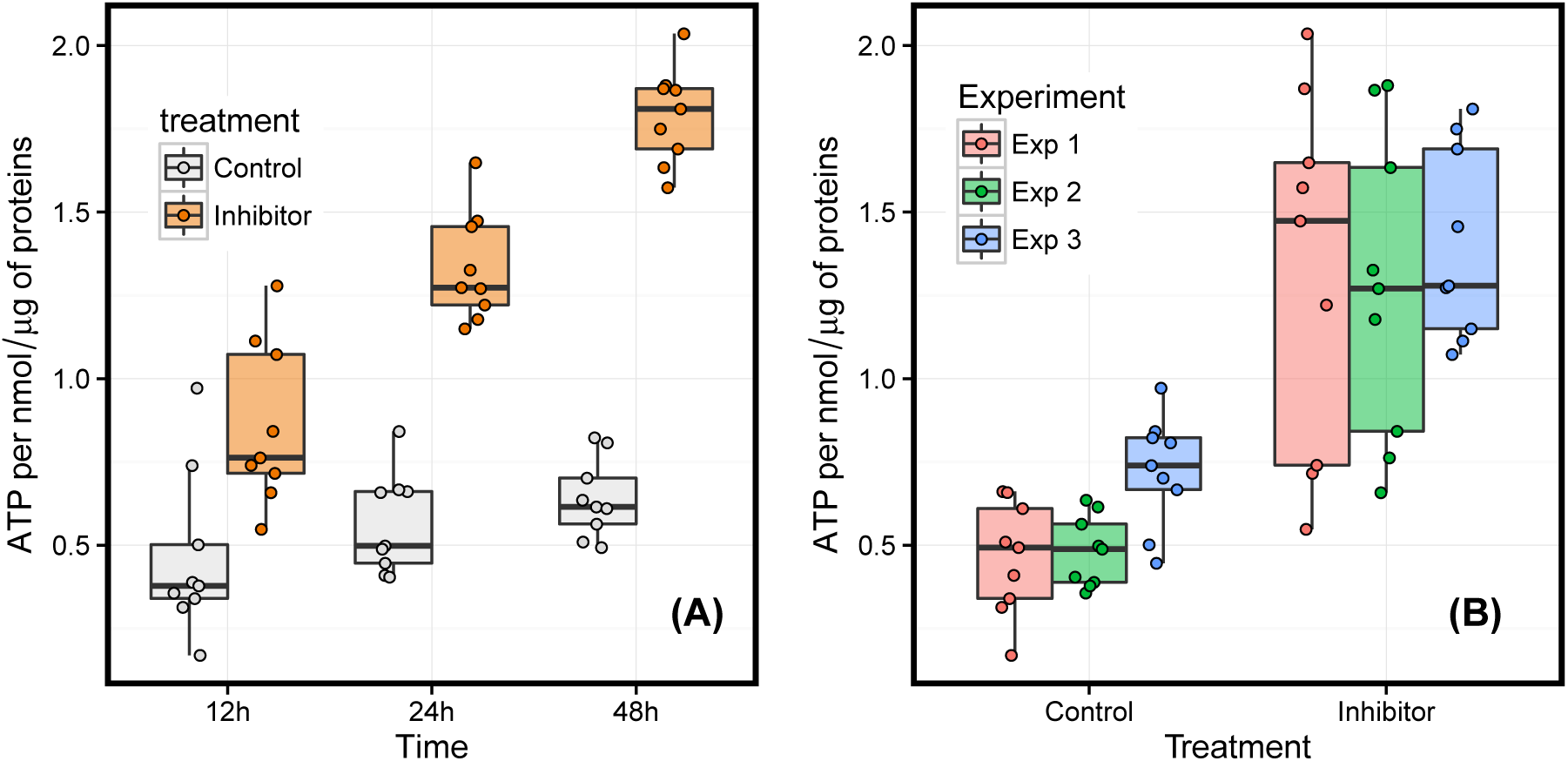
ATP production increases after inhibition of PknA/PknB. **(A)** ATP levels are higher for inhibitor treated samples compared to controls at all time points and **(B)** across three biological replicates (P = 3.213 × 10^*-*10^ for t-test comparing inhibitor vs control). Samples were normalized to the amount of residual protein (see Methods for details).

### Network clustering reveals condition-specific communities with different biological functions

While differential targeting analysis gives insight at the whole-network level, we aimed to understand biological organization at a more modular level. Because PANDA models groups of TFs regulating groups of genes, we naturally partitioned the nodes from each network into communities. To facilitate comparison of the network clusterings, we considered only the set of edges with positive edge weights (z-score greater than 0) in both conditions. This resulted in a Giant Connected Component (GCC) containing 67,740 edges between 143 TFs and 3971 genes. Genes that are also TFs were included as separate nodes in the network. As a network diagnostic, we plotted the distributions of edges per TF (outdegree) and edges per gene (indegree) (S1 Fig).

Next, we used CONDOR (COmplex Network Description Of Regulators), an R package for bipartite network analysis (*24*), to detect communities independently in each PANDA network. CONDOR maximizes the modularity, a score that can be interpreted as an enrichment for links within communities minus an expected enrichment given the network degree distribution.

This analysis identified 21 communities in the inhibitor network and 24 communities in the control network, with similar modularities of 0.495 and 0.504, respectively, and similar membership (S2 Fig). By testing each community as a whole for functional enrichment (see Methods), we found 4 of the 21 communities in the inhibitor network and 3 of the 24 communities in the control network to be functionally enriched (*FDR* < 0.05; overlap*>* 4) in one or more functional categories (S4 Table and S5 Table). This functional enrichment is robust to the choice of edge-weight threshold (Fig. S3). The differences in the enrichment identified between conditions are relevant in the context of PknA/PknB inhibition. For example, response to stress is enriched in a community in the inhibitor but not the control network; 11 of 26 genes in the functional category are assigned to community 2 in the inhibitor network, suggesting that these genes are cooperatively regulated in response to PknA/PknB inhibition. In addition to the activation of stress response, the results suggest that energy metabolism and ATP production regulation change in response to PknA/PknB inhibition.

To validate the network finding that ATP metabolism is disrupted by inhibition of PknA/PknB, we quantified the level of ATP in both the control and inhibitor treated cells at 12, 24, and 48 hours. After normalizing based on residual protein quantity (see Methods), we observed higher levels of ATP at all time points in the inhibitor treated samples compared to the control treatment (Fig. 4).

### Conclusion

As multi-omic data acquisition becomes increasingly commonplace, researchers interested in the drivers of complex phenotypes will face a new intellectual challenge: given a wealth of omics data, how can one identify relevant regulatory relationships amidst an abundance of correlations and partial correlations within and across data types? One solution to this challenge is to build regulatory networks that move beyond simple correlations by creating models that are consistent with known mechanisms of gene regulation. Here we applied one such algorithm, PANDA, to estimate the TF regulatory network response to an antimicrobial compound that inhibits two key signaling molecules important for cell wall function and stress response in *Mycobacterium tuberculosis*, PknA and PknB.

By comparing the regulatory networks from samples treated with either the PknA/PknB inhibitor or an inactive control compound, we identified treatment-specific network changes and provide validation for multiple network-generated hypotheses. This includes the observation that changes in TF outdegree correlated with changes in TF phosphorylation, suggesting that the PANDA regulatory networks successfully captured the relevant downstream effects of the perturbation. We identified “differentially targeting” PknA/PknB-specific TFs (based on change in TF outdegree) that affect diverse biological functions associated with signal transduction, cell wall function, and lipid metabolism. Additionally, we identified “differentially targeted” PknA/PknB-specific genes (based on change in gene indegree) that our models predict have a condition-specific pattern of regulation. In the case of mycobactin synthesis, these network changes are corroborated by changes in mycobactin lipid levels.

Networks are known to exhibit complex substructure, and we found the inferred regulatory networks are organized into “communities” of TFs and genes collectively associated with regulation of distinct biological processes. We used a bipartite community detection method to explore the structure of the *Mtb* regulatory network and found communities of genes and TFs that display condition-specific pathway enrichment. These include an inhibitor-specific community over-represented for genes involved in ATP production; this result was validated experimentally.

Taken together, these results demonstrate gene regulatory network inference using PANDA can effectively integrate multi-omic data and infer regulatory networks that capture downstream signaling effects. This, combined with advances in high-throughput methods for measuring phosphorylation-dependent protein-protein interactions (*25*), creates new opportunities for the functional characterization of drug candidates in *Mtb*. Approaches such as those described here will be essential for finding interventions in a disease that is already a substantial threat to human health and that is becoming increasingly difficult to treat.

## Acknowledgements

We thank Dr. Kimberly Glass for her assistance in interpreting the PANDA network results. This work was supported by grant R01AI099204 from the National Institute of Allergy and Infectious Diseases. JP is supported by the National Heart, Lung, and Blood Institute grant (K25HL140186), and JQ is supported by a grant from the National Cancer Institute (1R35CA220523).

## Materials and Methods

### RNA-seq and Phosphoproteomics Data

RNA-Seq and phosphoproteomics data collection, processing and results are available in (*6*). The RNA-Seq is available through the Gene Expression Omnibus (GEO) database under the accession number GSE110508, and protein phosphorylation data is available from the ProteomeXchange Consortium via the PRIDE partner repository with the dataset identifier PXD008968.

### Reconstructing PANDA networks

PANDA requires three inputs: a regulatory network prior of known TF-gene binding interactions, a protein cooperativity network prior of known protein-protein interactions, and expression data. We ran PANDA twice with the default update parameter (*α* = 0.10) using the same TF-gene and protein-protein interaction priors, but with gene expression data unique to samples treated with the active compound or control compound. Analyses were robust to different choices of the update parameter, *α*. PANDA outputs a final regulatory network, a final protein-cooperativity network, and a final co-expression network. All downstream analysis used only the regulatory network, which is represented by a collection of inferred TF-gene regulatory interactions.

### PANDA regulatory network prior

We created the regulatory network prior using ChIP-seq data from the supplemental material of (*7*). The regulatory network prior contains 6517 TF binding interactions between 143 TFs and 2501 genes, filtered by significance (*P* < 0.01) and located within the −150 to +70 nucleotide promoter window. In reconstructing networks, we consider only these 143 TFs as defined in (*7*) as potential regulators.

### PANDA Protein-cooperativity network prior

Predicted interactions between TFs were obtained from STRING v10 (*8*). We filtered these interactions to include only those between the 143 TFs in our regulatory network prior.

### ATP quantification

Triplicate samples from three individual experiments were grown and harvested at serial time points as previously described (*6*). Metabolic activity was rapidly quenched by placing the bacteria directly into 40% acetonitrile, 40% methanol, 20% water previously cooled on dry ice. The cells were then mechanically disrupted with 0.1 mm Zirconia beads in a MagNA Lyser instrument (Roche) by agitating the samples 4 times at 7000 rpm for 45 seconds with a cooling step at −20°C for 5 minutes between each cycle. The lysate was then clarified by centrifugation (10000g, 10 min, 4°C) and filtered through a 0.22 *µ*m filter (Costar^®^ #8160). ATP quantification was performed according to the manufacturer’s instructions of the ATP Colorimetric / Fluorometric Assay kit (BioVision #K354-100) and normalized to the residual protein quantity, determined by the Pierce^™^ BCA Protein Assay kit (ThermoFisher #23227).

### Functional annotations

Gene Ontology (GO) and KEGG Pathway functional categories were downloaded from PATRIC (*26*). We removed duplications and functional categories matching the regular expression `eukary|plant|(?i)photosynth|E.Coli|bile|insect`. To these we added manually curated functional categories (S6 Table).

### Statistical significance of PknA/PknB-specific TFs and genes

We used permutation testing to calculate empirical P-values for the significance of differential targeting for each TF and gene. To do this, we ran PANDA for each of 1000 randomized gene expression matrices with permuted gene labels. For each TF/gene, we then calculated significance by determining the proportion of the out/indegree differences for the 1000 runs that were greater the observed out/indegree if positive, or less than the observed out/indegree if negative.

### Bipartite network community detection

We used the CONDOR R package (*24*) with project=FALSE and other parameters set to default to detect communities in the inhibitor and control networks separately. These two networks contained the same subset of edges, but had different edge weights based on the output from running PANDA with data from the two conditions. Edges with weight < 0 in either network were removed from both networks.

### Functional enrichment analysis

To identify PknA/PknB-specific functional categories, we ran GSEA Preranked, which we downloaded from www.broadinstitute.org/gsea/ as the Java version. We ranked TFs/genes by their difference (inhibitor – control) in indegree/outdegree and ran GSEA using a minimum gene set size of 10 and a maximum size of 250. We report significant results (*FDR* < 0.1), with positive and negative enrichment scores representing enrichment in the inhibitor and control treatments, respectively. We used the one-sided Fisher’s Exact Test to evaluate the significance of each functional category for a given gene set. We required a minimum overlap of five genes between the gene set and the genes annotated to the functional category for significance to be considered. Multiple testing correction was done using the Benjamini-Hochberg method.

## Supporting Information

**S1 Fig**

**Degree distributions of thresholded PANDA networks.** (A) Histogram of the number of edges per transcription factor in each condition. (B) Histogram of the number of edges per gene in each condition.

**S2 Fig**

**Comparison of community overlaps between conditions.** The value in each cell is the number of (A) TFs or (B) genes assigned to both the corresponding inhibitor-treated and control-treated communities. The red shading helps to visualize how the TFs/genes in each control community are distributed among inhibitor communities.

**S3 Fig**

**The effect of threshold used to define PANDA subnetworks for CONDOR community detection.** (A) Number of edges that would be assigned to each subnetwork depending on the threshold (z-score cutoff) used. (B) Modularity of the network clustering per subnetwork depending on threshold. (C) Number of communities detected depending on threshold. (D) We select functional categories that show significant enrichment in at least 5 thresholds in either condition, together with functional categories that were significant at a threshold of 0. At each threshold and for each condition, we compute the proportion of genes in the network in the functional category present in the community containing the highest proportion of genes in the functional category. We then take the difference in this proportion between communities. For example, an inhibitor – control proportion of 0.5 may suggest that at a particular threshold, 0.75 of the genes in the functional category were present in the same community in the inhibitor, while only 0.25 of the genes in the functional category were present in the same community in the control network. We plot this difference in proportion for each functional category and at each threshold. Functional categories deemed significant at a threshold of 0 and discussed in the text are marked with an asterisk (*).

**S1 Table**

**Differential targeting of network TFs.**

**S2 Table**

**Differential indegree of network genes.**

**S3 Table**

**Differential targeting of functional categories identified by gene set enrichment analysis (GSEA).**

**S4 Table**

**Overrepresented functional categories in inhibitor-treated communities identified by the one-sided Fisher’s Exact Test.**

**S5 Table**

**Overrepresented functional categories in control-treated communities identified by the one-sided Fisher’s Exact Test.**

**S6 Table**

**Curated functional categories.**

